# Graph theoretical analysis reveals the adaptive role of the left ventral occipito-temporal cortex in the brain networks during speech processing

**DOI:** 10.1101/2022.02.03.478936

**Authors:** Shuai Wang, Samuel Planton, Valérie Chanoine, Julien Sein, Jean-Luc Anton, Bruno Nazarian, Anne-Sophie Dubarry, Christophe Pallier, Chotiga Pattamadilok

## Abstract

The left ventral occipito-temporal cortex (left-vOT) plays a key role in reading. Several studies have also reported its activation during speech processing, suggesting that it may play a role beyond written word recognition. Here, we adopt a graph theoretical analysis to investigate the functional role of this area in the whole-brain network while participants processed spoken sentences in different tasks. We find that its role and interactions with other areas changes in an adaptive manner. In a low-level speech perception task, the left-vOT is part of the visual network and acts as a connector that supports the communication with other cognitive systems. When speech comprehension is required, the area becomes a connector within the sensorimotor-auditory network typically recruited during speech processing. However, when comprehension is compromised due to degradation of speech input, the area disengages from the sensorimotor-auditory network. It becomes part of the visual network again and turns from connector into a simple peripheral node. These varying connectivity patterns are coherent with the Interactive Account considering the left-vOT as a convergent zone with multiple functions and interaction patterns that depend on task demands and the nature of sensory input.

## Introduction

Reading acquisition induces massive changes in brain functions, structures, and organization, especially within the auditory and visual systems (Dehaene et al., 2015). The most significant change is the emergence of an area, located in the left ventral occipito-temporal cortex (vOT), also known as the “Visual Word Form Area” (VWFA) due to its central role in reading (see Dehaene and Cohen, 2011 for a review). A number of studies have reported that once reading is acquired, this area consistently responds to known scripts, regardless of the characteristic of the writing system (Kuo et al., 2001; Hasson et al., 2002; Bolger et al., 2005; Nakamura et al., 2012; Szwed et al., 2014) and that the degree of activation is dependent upon individuals’ reading ability (Booth et al., 2003; Dehaene et al., 2010; Centanni et al., 2018; Kubota et al., 2019).

Interestingly, there is also empirical evidence that speech processing involves the left-vOT. Activation in response to speech input was found in various tasks, ranging from those that explicitly require a retrieval of spelling knowledge, such as determining whether spoken words share the same rime spelling (Booth et al., 2002; Cao et al., 2010) or contain a target letter (Ludersdorfer et al., 2015, 2016), to purely auditory tasks mimicking natural speech processing situations such as spoken word recognition or spoken sentence comprehension (Dehaene et al., 2010; Planton et al., 2019). The dominant explanation of this cross-modal activation has been proposed by Dehaene and Cohen’s research team: According to the *orthographic tuning hypothesis* (Dehaene and Cohen, 2011; Dehaene et al., 2015), left-vOT neurons are progressively tuned to written language input and become specialized in orthographic coding during reading acquisition. Nevertheless, these orthographic coding neurons could also be activated in a top-down fashion by spoken input. While this mechanism can account for the activation of the left-vOT during speech processing tasks that require the conversion of spoken inputs into the corresponding orthographic codes, or when a fast and automatic conversion can be conducted (Booth et al., 2002; Cao et al., 2010; Ludersdorfer et al., 2015, 2016), it remains unclear how it could explain the recruitment of the area in more ecological speech processing contexts where the online phonological-to-orthographic conversion seems unlikely (Planton et al., 2019). This observation raises a question about the general functional role of this area in the language network beyond its well-established contribution to reading.

An alternative view of the functional role of the left-vOT in widely distributed brain networks is the *Interactive Account* proposed by Price & Devlin, (2003, 2011; see also Buchel et al., 1998). According to the view, this area is an integration or a convergence zone that supports multiple functions depending on its interaction with other regions, and the nature of task and stimulus. As reviewed by Price and Devlin (2003), the same neural population within the area responded to non-orthographic inputs such as spoken words (Thompson-Schill et al., 1999; Price et al., 2003), objects (Moore et al., 1999; Phillips et al., 2002), colors (Price et al., 1997), and braille script (Buchel et al., 1998) and, therefore, seem to have multiple functions in different processing contexts depending on the interactions that they have with other cortical or subcortical areas. According to the authors, the prominent function of left-vOT in reading would arise from its ideal location at the transition between the occipital and the temporal lobe, enabling unique interactions between visual and language regions. Strong evidence supporting this view was provided by Saygin and colleagues (2016) in a longitudinal study in young children before (5-year-old) and after (8-year-old) they learned to read. The authors reported that the location of the area that became the VWFA in 8-year-old readers could be predicted from their pre-reading patterns of anatomical connectivity within the left occipitotemporal cortex, and that the “future VWFA” of 5-year-old children was preferentially connected with the spoken language areas. Studies conducted in adult readers also provide evidence that the left-vOT is connected with widely distributed regions through intrinsic connectivity (Koyama et al., 2010; Vogel et al., 2012; Stevens et al., 2017; Chen et al., 2019; Li et al., 2020) and anatomical connections (Yeatman et al., 2013; Bouhali et al., 2014; Chen et al., 2019).

In the case of speech processing, the Interactive Account considers that the activation of the left-vOT reflects functional integration between the phonological, orthographic and semantic representations. Within this theoretical framework, one can assume that the left-vOT acts as a connector that communicates with distributed brain regions and neural networks. The present study puts this hypothesis to test. We investigated the functional role of the left-vOT during different speech processing situations by modeling the brain as a network (Fedorenko et al., 2014; Hagoort et al., 2014). To this end, we applied an analysis based on graph theory to fMRI data from the study of Planton et al. (2019) collected while adult participants performed either low-level perception or comprehension task on spoken sentences. Within each task, spoken sentences were presented either against a silent background or against unintelligible “multi-speaker” babble noise, mimicking a “cocktail party” situation. Planton et al (2019) reported that the two manipulations had significant impacts both on performance and brain activation across different brain areas, including the left-vOT.

Graph theory provides a quantitative tool to investigate the organization of brain networks and interactions between brain regions (Bullmore et al., 2009; Rubinov et al., 2010). In the present study, the brain networks were constructed as a graph consisting of nodes and edges, in which nodes are regions of interest (ROIs) and edges are functional connectivity between ROIs. The organization of brain network and the functional role of the left-vOT within the network were examined in different speech processing conditions which included a “baseline” condition where participants were passively exposed to multi-speaker babble noise and “active” conditions where participants conducted the two sentence processing tasks in the two types of background noise as described above. The present study carried out the graph theoretical analysis of brain networks at three levels. First: *The global level* where the organization of the whole brain network is described by global measures of integration (*global efficiency)* and segregation (*clustering coefficient*). These two measures respectively describe the efficiency of information exchange across the whole brain and within nearest neighbors of nodes. Second: *The sub-network (community) level*. Here the global network is subdivided into different sub-networks which often correspond to specialized functional components (Power et al., 2011; Sporns et al., 2016). This level of analysis allowed us to identify the sub-network to which the left-vOT belonged in the different tasks. Third: *The nodal level*. Here, we specified, for each speech processing condition, the functional role of the left-vOT within the network, using different measures described in details in the Method section (Network Analysis), to examine: (1) whether the left-vOT acted as a connector as postulated by the Interactive Account, and (2), whether its functional role varied with the task and context (silence vs. babble noise) and, (3) how it correlates with task performance. With regard to the first question, the *participation coefficient* is of particular interest, which indicates whether the left-vOT acts as a connector in a network (Guimerà et al., 2005; van den Heuvel et al., 2013), on the basis of its connections with distributed brain regions and sub-networks. The second and third questions allowed us to assess whether the functional role of this area is dependent on the nature of the processing context and task difficulty.

## Methods

### Participants

Twenty-four native French speakers were recruited in the study (mean age: 24.05±3.46, 11 females). Participants were healthy, right-handed, with normal hearing and vision and reported no past or current neurological or language disorders. Written informed consents were obtained from all participants. The study was approved by the local ethics committee (CPP Sud Méditerranée #RCB 2015-A00845-44).

### Tasks and Stimuli

#### Spoken sentence processing tasks

The stimuli were spoken sentences expressing true or false statements. Both tasks used a Go/NoGo paradigm. In the low-level perception task, participants were instructed to press the response button as soon as they heard the same sentence twice in a row (Go trials). In the comprehension task, they had to press the response button whenever they heard a false statement, thus requiring a complex semantic analysis (Go trials). For both tasks, all NoGo trials were true statements. The two tasks were alternately presented in four separate runs (two runs per task). The run order was counterbalanced across participants. Each run lasted 7.2 min. and contained 10 Go trials and 70 NoGo trials that were pseudo-randomly distributed into 20 blocks of 4 trials. Two consecutive go trials were avoided. In half of the blocks, the spoken sentences were presented against clear background and half were presented against unintelligible multi-speaker babble noise at an SNR of +6 dB. In each participant, each NoGo sentence was presented only once but across participants, it appeared equally in the four active listening conditions (perception of clear speech, PN-; perception of speech-in-noise, PN+; comprehension of clear speech, CN-; comprehension of speech-in-noise, CN+). In addition to these “active” blocks, 5 “rest” blocks corresponding to silent background and 5 “rest” blocks corresponding to the multi-speaker babble noise were added to the run. Each of the active and rest blocks lasted 14s on average (range 12s–18s). Within each run, the order of the blocks from the different conditions was pseudorandomized to avoid two consecutive blocks of the same condition. At the trial level, each spoken sentence lasted from 1s-2.4s. During this period, a visual fixation cross was presented on the screen. After the sentence presentation, there was a blank screen whose duration was jittered. The SOA (3.55s on average) followed an exponential curve to maximize design efficiency (see Henson, 2015). The same procedure was adopted during the rest trials, except that the sentence was replaced by silence or multi-speaker babble noise.

#### Visual localizer task

To individually localize the left-vOT, the participants performed a functional localizer task in which sequences of 6-letter mono or disyllabic words and 6-letter consonant strings were visually presented and participants were required to detect, by pressing the response button, twelve target stimuli (Go trials: “######”) that were randomly included in the sequences. The task was presented in a single run that lasted 7.4 min. During the run, words and consonant strings were grouped in short blocks of ∼12s each (range 11s–13.3s). Each block contained 24 stimuli of the same category. Altogether, there were 12 word-blocks and 12 consonant-string blocks. For both categories, each stimulus remained on the screen for 340 ms and was followed by a blank screen of variable duration (∼160 ms on average). In addition to these 24 “active” blocks, 12 “fixation” blocks during which a cross remained on the screen throughout the block duration (∼ 12s on average) were also included. The 36 blocks were presented in a pseudorandom order to avoid repetition of the same condition. For all stimulus types, the visual input always appeared in the center of the screen, in white font on a dark grey background. Word stimuli were nouns and adjectives selected from the French database LEXIQUE (lexical frequency ∼7.21 per million, on average, New et al., 2004). No words or consonant strings were presented twice during the run. The analysis conducted in the visual localizer task is presented in the SI, Methods A.

### fMRI Data Acquisition and Pre-processing

The experiment was conducted on a 3T Siemens Prisma Scanner (Siemens, Erlangen, Germany) at the Marseille MRI center (Centre IRM-INT@CERIMED, UMR7289 CNRS & AMU, http://irmf.int.univ-amu.fr/) using a 64-channel head coil. T1-weighted images were acquired using an MPRAGE sequence (voxel size = 1×1×1 mm^3^, data matrix = 256×256×192, TR/TI/TE = 2300/900/2.98 ms, flip angle = 9º). Fieldmap images were also obtained using Dual echo Gradient-echo acquisition (TR = 677 ms, TE1/TE2 = 4.92/7.38 ms, FOV = 210×210 mm^2^, voxel size = 2.2×2.2×2.5 mm^3^). Functional images were collected using a gradient EPI sequence (TR = 1224 ms, TE = 30 ms, 54 slices with a thickness of 2.5 mm, FOV = 210×210 mm^2^, matrix = 84×84, flip angle = 66º, multiband factor = 3). Auditory hardware channel was composed of the Sensimetrics S14 MR-compatible insert earphones with a Yamaha P-2075 power amplifier.

Pre-processing was conducted by using fMRIPrep 20.0.6 (Esteban et al., 2019). For more details on the preprocessing pipeline, see fMRIPrep’s documentation (https://fmriprep.org/en/20.0.6/workflows.html). The T1-weighted image was corrected for intensity non-uniformity with N4BiasFieldCorrection in ANTs (Avants et al., 2008; Tustison et al., 2010), and used as T1w-reference throughout the workflow. The T1w-reference was then skull-stripped. The brain-extracted T1w was used for segmentation of cerebrospinal fluid (CSF), white-matter (WM) and gray-matter (GM) using fast (FSL 5.0.9). Volume-based spatial normalization to the standard MNI space was performed through nonlinear registration with antsRegistration, using brain-extracted versions of both T1w reference and the T1w template (MNI152NLin2009cAsym). For functional images, the fieldmap distortion correction was performed based on a phase-difference map. The functional images were then co-registered to the T1w reference using flirt (FSL 5.0.9) with the boundary-based registration (Greve et al., 2009) with nine degrees of freedom. Head-motion parameters were estimated before any spatiotemporal filtering using mcflirt (FSL 5.0.9). Fieldmap distortion correction, head-motion correction, BOLD-to-T1w co-registration, and spatial normalization were carried out in a single interpolation step by composing all the pertinent transformations. The pre-processed BOLD data were then used to calculate several confounding time series, including framewise displacement (FD), the mean signals within the white matter and the CSF, and a set of principal components of white matter and CSF that were extracted by the aCompCor method (Behzadi et al. 2007).

### Network Construction

The present study aimed to investigate the functional role of the left-vOT in brain networks during speech processing. In order to construct brain networks, a set of 264 regions of interest (ROIs) was taken from Power et al. (2011). Each ROI is a sphere with a radius of 5 mm and contains 81 voxels. The 264 spherical ROIs covers the entire cerebral cortex, subcortical areas and the cerebellum. This set of ROIs were intersected with the group-averaged gray matter mask to exclude areas that are outside the gray matter. One ROI at right thalamus (MNI x = 9, y = -4, z = 6) was removed due to no overlap with the group-averaged gray matter, resulting in a set of 263 ROIs. In addition to these ROIs, the left-vOT identified at the group level in the visual localizer task (see **Tasks and Stimuli** and **SI Methods A**) was included (centre MNI x = -47, y = -55, z = -17). To combine the left-vOT with the set of 263 ROIs, the group left-vOT was extracted with a volume of 81 voxels by adjusting the significance threshold (T > 5.2536, *p* < 2.5e-5). One ROI at left Fusiform gyrus (MNI x = -47, y = -51, z = -21) was further removed because it overlapped with the group left-vOT, thus resulting in a final set of 263 ROIs as the nodes in the networks (see **SI Fig. S1A** for the spatial distribution of the set of ROIs).

Pre-processed functional data were scaled to percent of signal change and modeled by using the Least Squares — Separate (LSS) method (Mumford et al., 2012; 3dLSS in AFNI) which ran a GLM for each trial and output trial-wise estimates (i.e., β coefficients) for each condition. To estimate the edges of the networks in the baseline (i.e., noise-only) and active sentence listening conditions, we used a beta-series connectivity analysis (Rissman et al., 2004), which estimates functional connectivity between two regions by calculating the correlation of the activity (i.e., beta estimates) of the two regions across trials. Specifically, the six motion parameters, their temporal derivatives, and all their corresponding squared time series (i.e., 24 head motion regressors) were included in the LSS models to control for the impacts of head motion (Friston et al., 1996). In addition, the mean time-series and the first twelve principal components of white matter and of CSF were extracted by using the aCompCor method (Behzadi et al., 2007) and used as nuisance regressors in the LSS models to reduce influence of physiological noise. The cosine-basis regressors estimated by fMRIPrep for high-pass filtering were also included in the LSS models as nuisance regressors. Motion contaminated volumes were identified by using framewise displacement (FD) and were censored along with the prior volume if their FD > 0.5mm. On average, 2.2% of the volumes were censored. The trial-wise beta estimates were then used to calculate beta-series connectivity. The series of beta estimates were averaged over voxels within each ROI for each condition. To estimate beta-series connectivity, Fisher-z-transformed Spearman correlation of averaged beta series between each pair of ROIs was calculated, resulting in an undirected 263×263 correlation matrix for each condition per participant. Each correlation matrix was thresholded into a binary matrix at a target density x% by keeping the x% of strongest edges to construct a brain network. The range of density between 15% and 22% (at intervals of 1%) was selected based on the largest connected component (LCC), which was calculated to ensure that the networks are largely connected since networks tend to be unstable and contain more fragments at lower densities, while they become more random at higher densities (Bullmore et al., 2011). The lower bound 15% density is the sparsest density at which 90% of all the networks are fully-connected (i.e., the size of the LCC equals the size of the network), without any significant between-condition difference in the size of the LCC. The upper bound 22% density is the sparsest density where all the networks are fully-connected. The results at the lower bound 15% were reported in the main text. The Supplementary Information (**SI Results**) reported the results across the range of densities (15%-22%) with the false discovery rate (FDR) correction to ensure the results did not rely on a single density.

### Network Analysis

Focusing on the left-vOT node, the graph theoretical analysis was carried out to characterize three topological scales of the brain network, i.e., from global to sub-network and nodal level (see **Table 1** for the interpretations of graph measures). The analysis was applied to each participant and each condition.

**Table 1.**
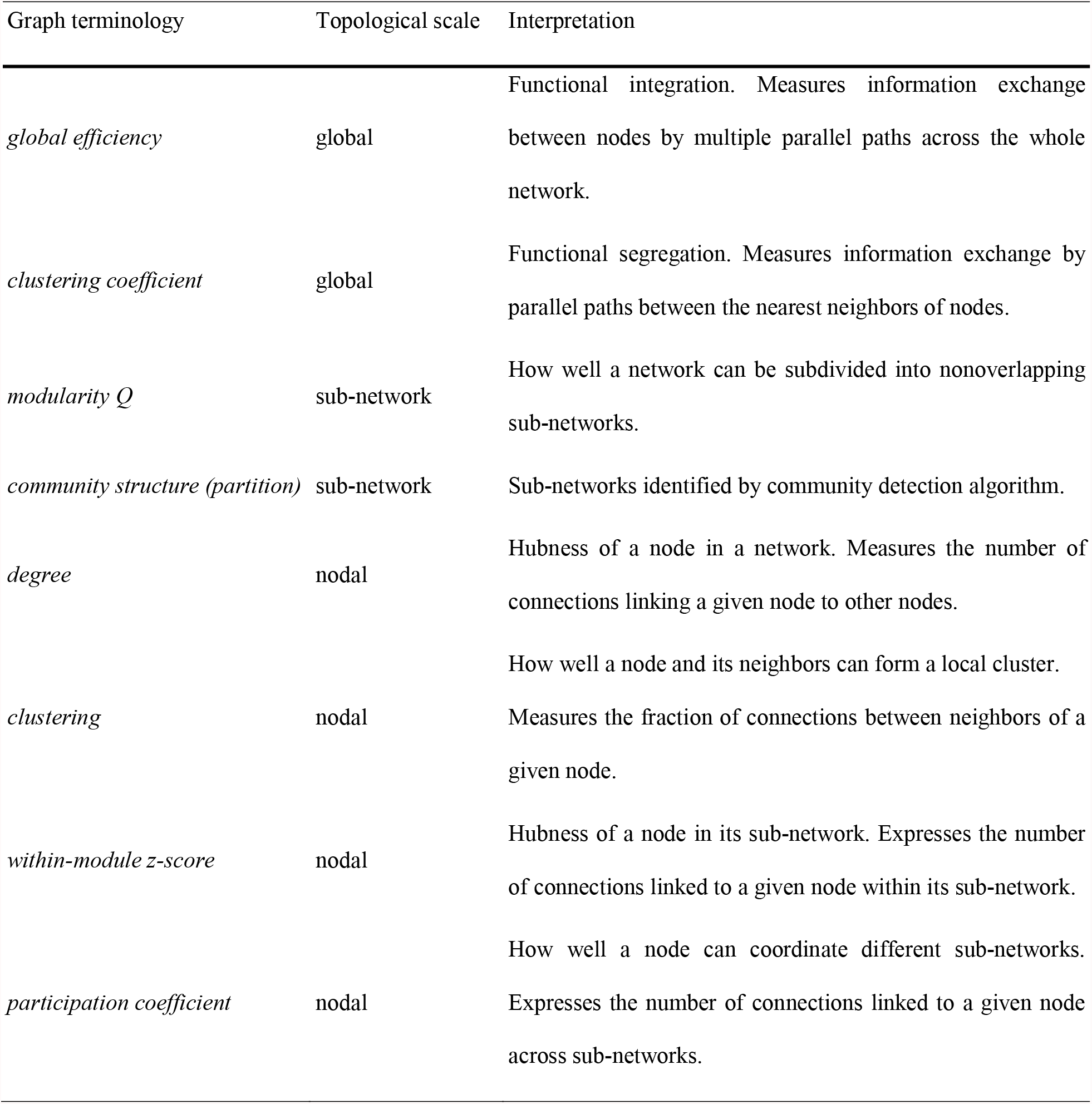
Interpretations of graph theory terms.

| Graph terminology | Topological scale | Interpretation |
| --- | --- | --- |
| *global efficiency* | global | Functional integration. Measures information exchange between nodes by multiple parallel paths across the whole network. |
| *clustering coefficient* | global | Functional segregation. Measures information exchange by parallel paths between the nearest neighbors of nodes. |
| *modularity $Q$* | sub-network | How well a network can be subdivided into nonoverlapping sub-networks. |
| *community structure (partition)* | sub-network | Sub-networks identified by community detection algorithm. |
| *degree* | nodal | Hubness of a node in a network. Measures the number of connections linking a given node to other nodes. |
| *clustering* | nodal | How well a node and its neighbors can form a local cluster.<br>Measures the fraction of connections between neighbors of a given node. |
| *within-module z-score* | nodal | Hubness of a node in its sub-network. Expresses the number of connections linked to a given node within its sub-network. |
| *participation coefficient* | nodal | How well a node can coordinate different sub-networks.<br>Expresses the number of connections linked to a given node across sub-networks. |

Firstly, to examine the global changes induced by active sentence processing compared to baseline, two global graph measures, i.e., *global efficiency* and *clustering coefficient*, were estimated to characterize the functional integration and segregation of the whole-brain network, respectively. *Global efficiency* is a measure of the capacity of the network for global information transfer. In other words, it measures information exchange between nodes by multiple parallel paths across the whole network. *Clustering coefficient* is a measure of the local efficiency of information transfer by parallel paths between the nearest neighbors of nodes (Bullmore et al., 2009).

Secondly, to identify the sub-network to which the left-vOT belonged in the different speech processing contexts, community detection was conducted to subdivide the whole brain network into several sub-networks by relatively maximizing intra-connections and minimizing inter-connections. For each participant and each condition, the Louvain algorithm (Blondel et al. 2008) and the consensus partitioning (Lancichinetti et al., 2012) were applied on the whole brain network to determine the optimal partition, the corresponding *number of communities*, and *modularity Q*. Specifically, the Louvain algorithm (*γ* = 1) was performed 1,000 times on the network to generate 1,000 initial optimal partitions that were used to estimate the agreement matrix, which was then submitted into the consensus partitioning (□ = 0.5, N_iter_ = 1,000) to converge to an optimal partition. The optimal partitions from the different participants were further grouped and submitted into the consensus partitioning again to converge to a single representative partition for each condition. The representative partition of each condition was used to identify all the sub-networks and the one to which the left-vOT node belonged. The anatomical locations of the sub-networks were visually inspected and labelled, by comparing them with the community structures defined by Power et al. (2011).

Lastly, the nodal topology was estimated for each participant and each condition to characterize the functional role of the left-vOT node in the whole brain network during speech processing. Four nodal measures, including *degree, clustering, within-module z-score*, and *participation coefficient* were estimated. *Degree* or *degree centrality* measures the number of connections linking a given node to other nodes. Node with high *degree*, named high-degree hub, plays as a hub in information exchange by directly interacting with numerous nodes in the whole brain network. *Clustering (nodal)* measures the fraction of connections between neighbors of a given node. Node with high *clustering* is a part of a densely interconnected local cluster that contributes to the functional segregation of a network. Based on the community structure, *within-module z-score* and *participation coefficient* were estimated (Guimerà et al., 2005). *Within-module z-score*, as a sub-network version of *degree*, expresses the number of connections linked to a given node within its sub-network. Node with high *within-module z-score* is a high-degree hub in its own sub-network. *Participation coefficient* expresses the number of connections linked to a given node across different sub-networks. Node with high *participation coefficient*, that is a connector, plays a central role in coordinating the communications between different sub-networks. The network analysis was carried out by using the Brain Connectivity Toolbox (Rubinov et al., 2010). To assess the significance of these graph measures, the repeated measures permutation test (Asymptotic General Independence Test) was adopted to compare multiple conditions, and the pairwise permutation test was used as a post-hoc test. The statistical tests were conducted by using R, the “coin” package (Hothorn et al., 2006) and the “rcompanion” package (Mangiafico 2020). In addition, to confirm the main results, the analyses described above were conducted again by using a symmetrical set of ROIs (Di et al., 2014; for details, see **SI Methods B and Fig. S1B**).

## Results

### Global network changes induced by speech processing

To illustrate the overall organization of the brain network, as well as the pattern of the connections of the left-vOT node at the functional state where unintelligible speech input was presented, the individual baseline networks were accumulated across participants (**Fig. 1A**). The accumulated baseline network reflected a modular organization (see the matrix in **Fig. 1A**), where the left-vOT node was connected with widely distributed brain regions (see the glass brain in **Fig. 1A**).

**Figure 1.**
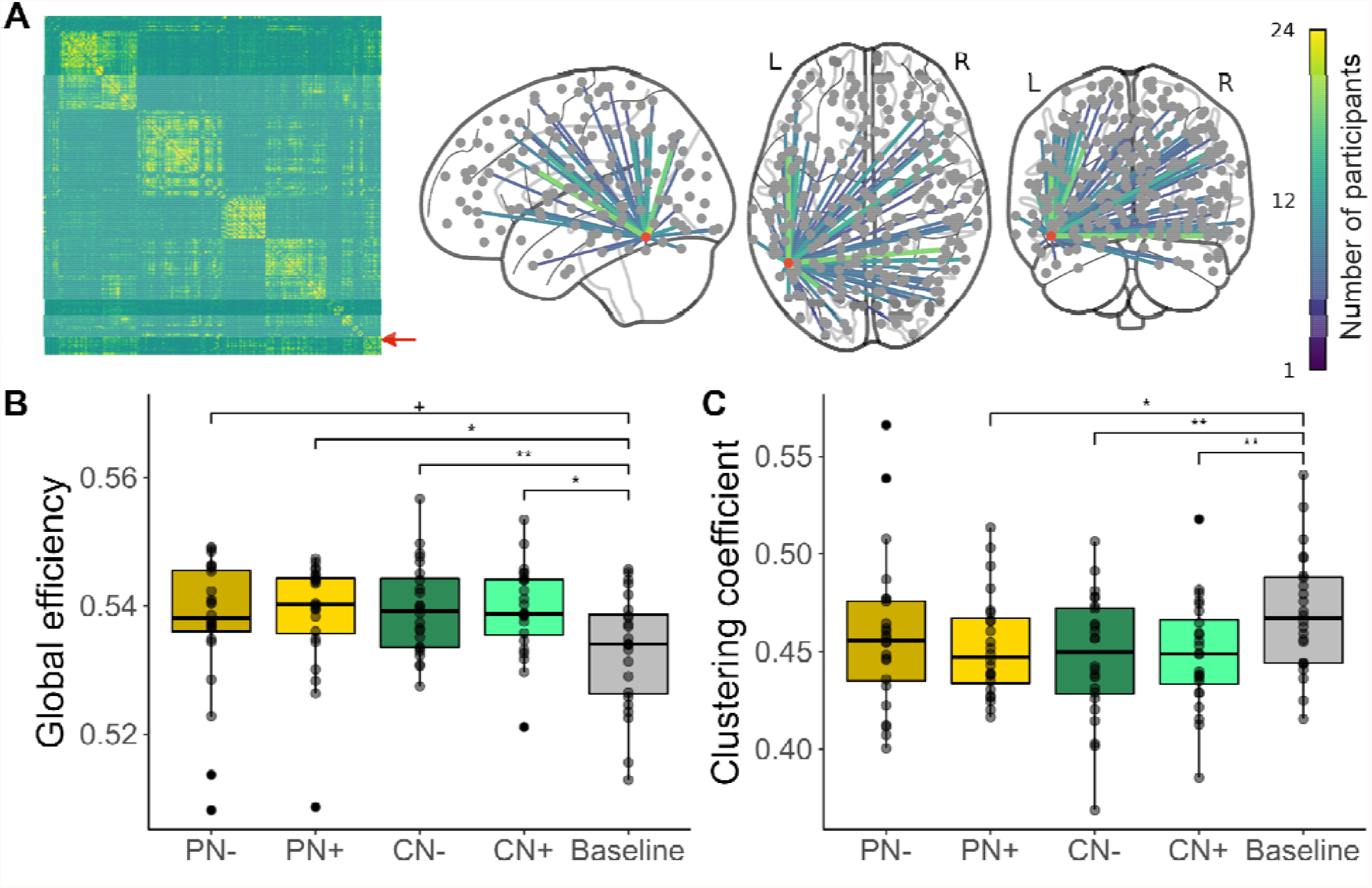
Overall network organization and global topology. **A**. The network of the baseline condition was used as reference to illustrate the overall network organization. The accumulated baseline network is shown as a matrix (left panel). The connections of the left-vOT node (indicated by the red arrow) are extracted from the accumulated network and shown on the glass brain (right panel, the red dot represents the left-vOT node and only connections presented in more than 25% participants are shown). **B**. The speech processing conditions have higher *global efficiency* than the baseline condition. **C**. The speech processing conditions have lower *clustering coefficient* than the baseline condition. **: p < 0.01; *: p < 0.05; + (marginal): p < 0.06. PN-: perception clear speech; PN+: perception speech-in-noise; CN-: comprehension clear speech; CN+: comprehension speech-in-noise.

Firstly, we examined whether processing speech in different experimental conditions induced network changes in global topology compared to the baseline network. Significant global network changes were found in both *global efficiency* and *clustering coefficient* (**Fig. 1 B and C**). Compared to the baseline network, the networks of the speech processing conditions showed an increase in *global efficiency* (*p* < 0.0022; see **Fig. 1B**). The *post-hoc* pairwise comparisons showed that the *global efficiency* of the PN+, CN- and CN+ conditions are significantly higher than the baseline condition (all *post-hoc p*s < 0.016), while the difference between the PN- and the baseline is marginally significant (*post-hoc p* < 0.056). No significant differences were found between the four speech processing conditions. These results indicate an overall higher level of communication between brain regions during speech processing in comparison to the baseline, i.e., when participants were presented to unintelligible multi-talker babble noises without task demand.

Moreover, in contrast to the baseline condition, a reduction of *clustering coefficient* was observed in the speech processing conditions (*p* < 0.0059; **Fig. 1C**) where the significant increases of global efficiency were observed (PN+: *post-hoc p* < 0.025; CN-: *post-hoc p* < 0.0074; CN+: *post-hoc p* < 0.0078) suggesting that the increase in global efficiency was accompanied by reduced interconnections between topological local neighbors.

Overall, the results of global network changes suggest that speech processing leads to a more integrated and less segregated network. The fact that the networks of the four speech processing conditions did not differ significantly in terms of global topology allows us to directly compare the community structures and nodal topology between them.

### Left-vOT node switching between sub-networks

In this part, the community structures were identified for each condition. Then, we examined the sub-network switching of the left-vOT node between two conditions. Based on the community structures defined by Power et al. (2011), four sub-networks were identified for both baseline and speech processing conditions (**Fig. 2**): visual network, fronto-parietal network, default mode network, and sensorimotor-auditory network (see **SI Methods C and Fig. S2** for details on sub-network labeling). The community structures are similar across conditions as shown by the normalized mutual information (NMI) where all similarities were higher than 0.73 (the range of NMI is [0, 1]; see the left-bottom panel in **Fig. 2**). This observation was also confirmed by the absence of significant difference across conditions in both modularity Q (*p* > 0.39) and number of communities (*p* > 0.097). The flow diagrams in **Fig. 2** illustrating the reconfigurations of sub-networks across conditions showed that the reconfigurations only involve a limited number of brain regions (the width of flow indicates the number of nodes).

**Figure 2.**
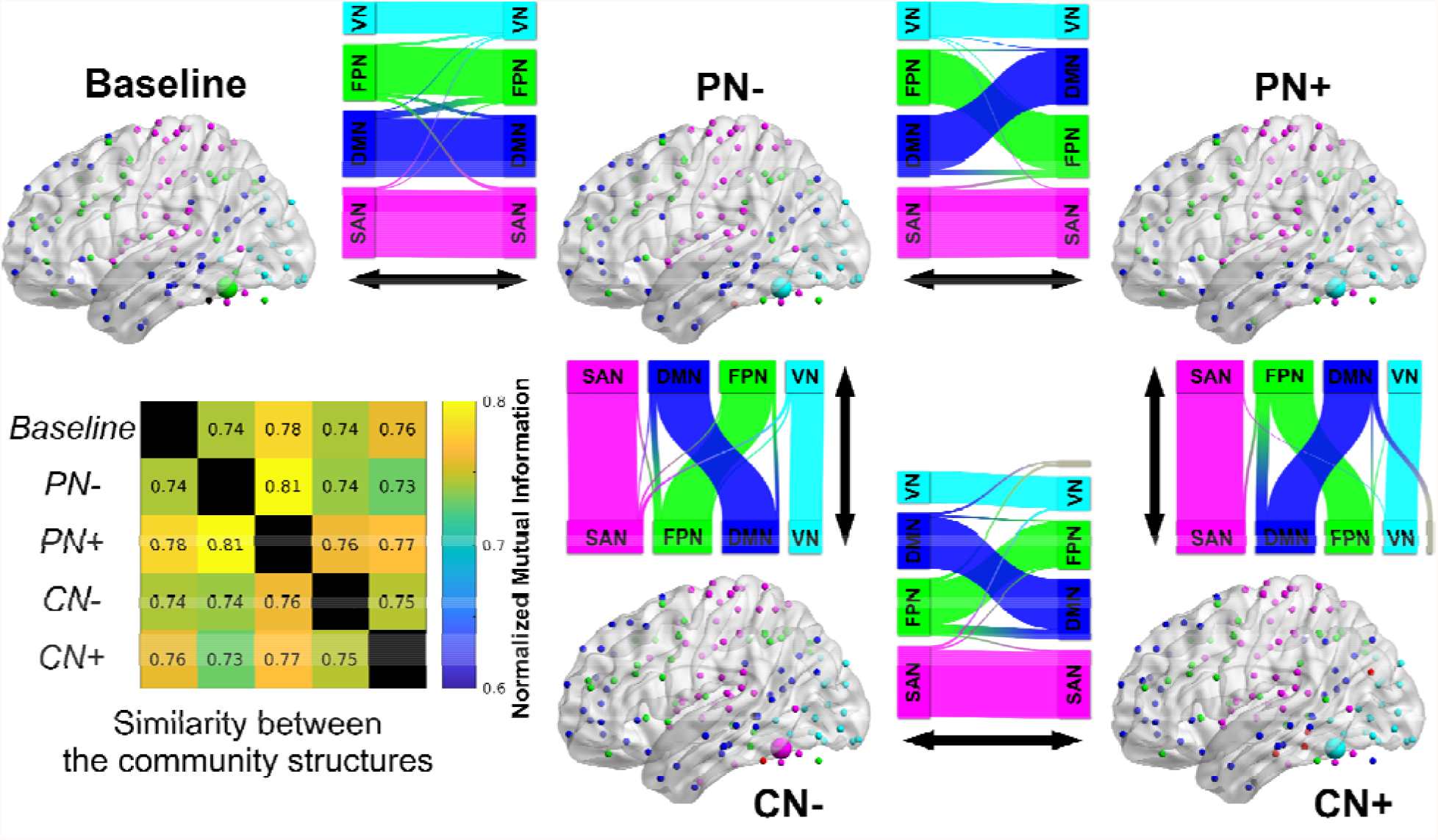
Community structures and switching of the left-vOT node across conditions. The baseline network and the four speech processing networks have similar community structures, which consist of visual network (VN, colored cyan), fronto-parietal network (FPN, colored green), default mode network (DMN, colored dark blue), and sensorimotor-auditory network (SAN, colored magenta). The matrix located in the left-bottom panel shows the similarity (normalized mutual information, NMI) between the community structures of different conditions. The brain maps illustrate that the left-vOT node (the largest bubble in each brain map, whose color refers to the corresponding sub-networks) belonged to different sub-networks in different conditions. The flow diagrams illustrate the reconfiguration of sub-networks across conditions (the width of the flows indicates the number of nodes). The brain maps were visualized with the BrainNet Viewer (Xia et al., 2013, http://www.nitrc.org/projects/bnv/). The flow diagrams were visualized with the MapEquation software (Rosvall et al., 2010).

More specifically related to the present research question, analyses focusing on the left-vOT node showed that the node (illustrated by the largest bubble in each brain map presented in Figure 2) participates in the reconfiguration of the sub-networks as it was identified as a part of different sub-networks in different conditions. Specifically, at the initial stage (in the baseline condition), the left-vOT was a part of the fronto-parietal network. When the participants performed the perceptual task without noise (PN-), the area became part of the visual network, and it remained in this network even when the background noise was added (PN+). Interestingly, the area switched from the visual network to the sensorimotor-auditory when the participants had to extract the semantic content of clear sentences in the comprehension task (CN-). Finally, it returned to the visual network when the background noise was added during the comprehension task (CN+).

### Nodal topology of the left-vOT node

Next, four nodal measures, including *degree, clustering, within-module z-score*, and *participation coefficient*, were estimated to characterize the functional role of the left-vOT node in brain networks. For each nodal measure, the group averages were calculated and were used to rank all of the 263 nodes considered in the analysis. In order of *degree*, the left-vOT node was ranked 66^th^ (PN-), 18^th^ (PN+), 142^th^ (CN-), and 50^th^ (CN+) among the 263 nodes. In order of *clustering*, it was ranked 259^th^ (PN-), 248^th^ (PN+), 256^th^ (CN-), and 249^th^ (CN+). Neither *degree* nor *clustering* showed significant differences between the conditions. These results suggest that the left-vOT is unlikely to be a degree-based hub and, with neighbors that likely disconnect with each other (i.e., low *clustering*), it tends to connect with widely distributed regions rather than joining a local cluster.

Based on the partitions of sub-networks described above, *participation coefficient* and *within-module z-score* were examined together to define the functional role of the left-vOT, i.e., whether it acted as a connector. In PN-, PN+ and CN-conditions, the left-vOT node was ranked among the top 3 nodes with highest *participation coefficient*, whereas its rank in the CN+ condition was dropped to 40^th^ (**Fig. 3A)**. The ranks of the left-vOT node on *within-module z-score* ranged between 75^th^ and 188^th^ across the four conditions (**Fig. 3B)**. The comparison between the four speech processing conditions revealed a significant difference in *participation coefficient* (*p* < 0.0012; **Fig. 3C**). The *post-hoc* tests confirmed that the *participation coefficient* in the CN+ condition was significantly lower than the PN-(*p* < 0.0059), the PN+ (*p* < 0.0034), and the CN-(*p* < 0.028) conditions. The comparison in *within-module z-score* did not find a significant difference (*p* > 0.075). Taken together, the high *participation coefficient* and intermediate *within-module z-score* observed in the PN-, PN+ and CN-conditions indicate that the left-vOT node acts as a connector in the network in these processing contexts. In the CN+ condition, where the task performance was also the poorest (cf. Planton et al., 2019) both in terms of the accuracy scores (PN-: 96%, PN+: 92%, CN-: 88%, CN+: 63%) and the reaction times (PN-: 1296 ms, PN+: 1413 ms, CN-: 2389 ms, CN+: 2575 ms), both *participation coefficient* and *within-module z-score* were at the intermediate level, suggesting that the left-vOT behaves like a peripheral node in the network. Altogether, these results suggest that the communications between the left-vOT node and other sub-networks are reduced in the CN+ condition, while in the other conditions the left-vOT node holds the highest level of communication with other sub-networks across the whole brain network.

**Figure 3.**
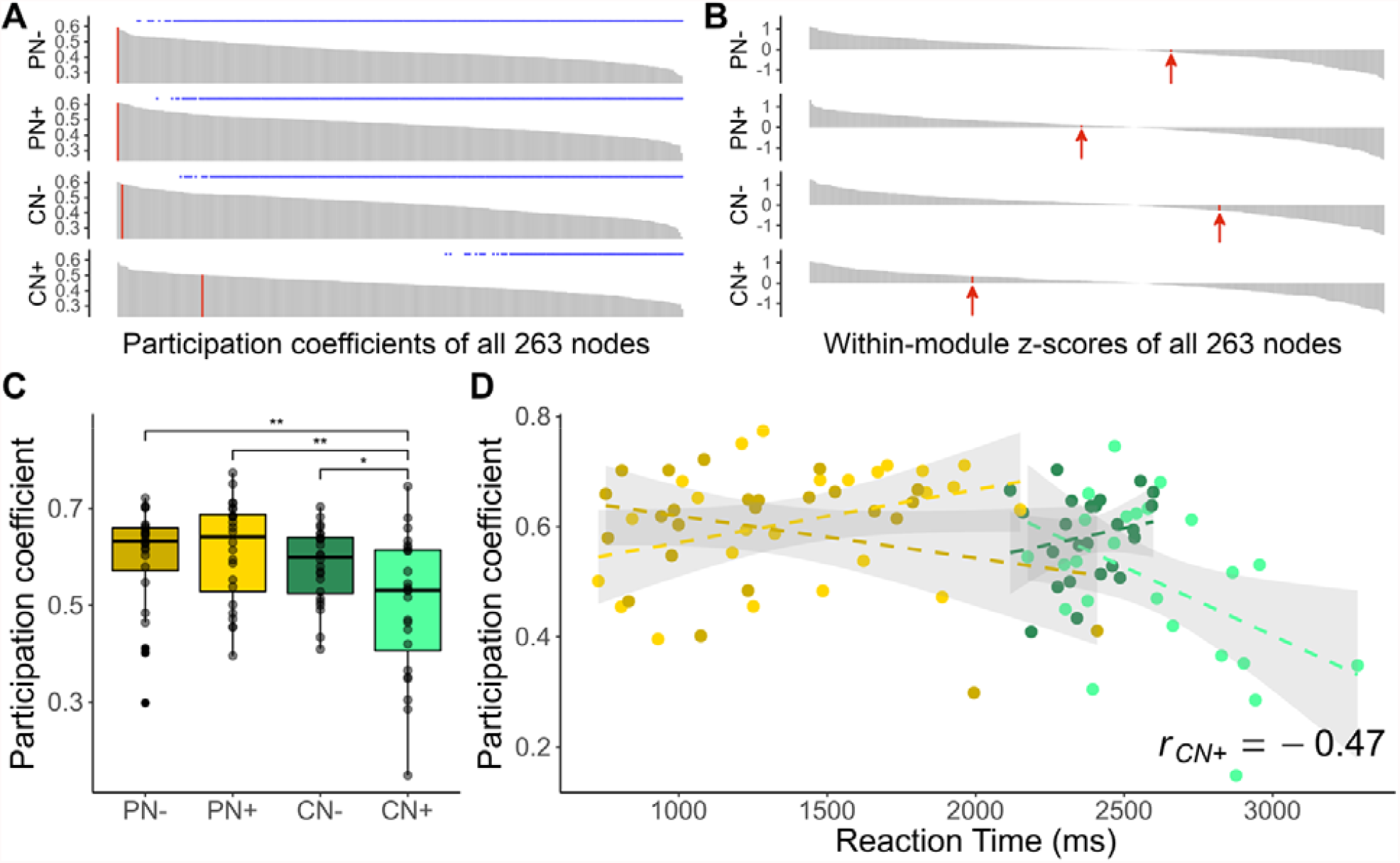
Nodal topology of the left-vOT node and its relationships with reaction time. **A**. Across the four speech processing conditions, the *participation coefficient* of the left-vOT node was ranked 1^st^ (PN-), 1^st^ (PN+), 3^rd^ (CN-), and 40^th^ (CN+) among the 263 nodes. The vertical red bar indicates the rank of the left-vOT node among the 263 nodes considered in the analysis. The blue asterisks indicate the nodes that showed significantly lower *participation coefficients* than the left-vOT (paired permutation test, *p* < 0.05 unc.). **B**. Across the four speech conditions, the *within-module z-score* of the left-vOT node was ranked 166^th^ (PN-), 125^th^ (PN+), 188^th^ (CN-), and 75^th^ (CN+) among the 263 nodes. The vertical red bars (also indicated by the red arrows) indicate the left-vOT node. **C**. The *participation coefficient* of the left-vOT node was significantly lower in the CN+ condition than in the PN-, PN+ and CN-conditions. **D**. The *participation coefficient* and reaction time was negatively correlated in the CN+ condition (light green dots and dashed line; Pearson’s r = -0.47, *p* < 0.022), but not in the other conditions (all *p*s > 0.078).

Having identified the functional role of the left-vOT as a connector in the PN-, PN+, and CN-conditions, we further examined the relationship between the values of *participation coefficient* of the left-vOT and the task performance to investigate the contribution of the area to speech processing. As shown in **Fig. 3D**, the correlation analysis showed that *participation coefficient* and reaction time was negatively correlated in the CN+ condition (Pearson’s r = - 0.47, *p* < 0.022), but not in the other conditions (all *p*s > 0.078). This indicates that participants who responded more quickly in the speech comprehension task that was conducted in a noisy environment were those who showed more connections between the left-vOT and the different sub-networks.

The nodal results of the left-vOT reported here were further validated in supplementary analyses using a symmetrical set of ROIs (Di et al., 2014; for details, see **SI Results and Fig. S3**).

## Discussion

The present study investigated the functional role of the left-vOT in speech processing by applying graph theoretical analysis to fMRI data collected during spoken sentence listening in different tasks (Planton et al., 2019). **Figure 4** summarizes the main observations obtained at the three levels of analysis.

**Figure 4.**
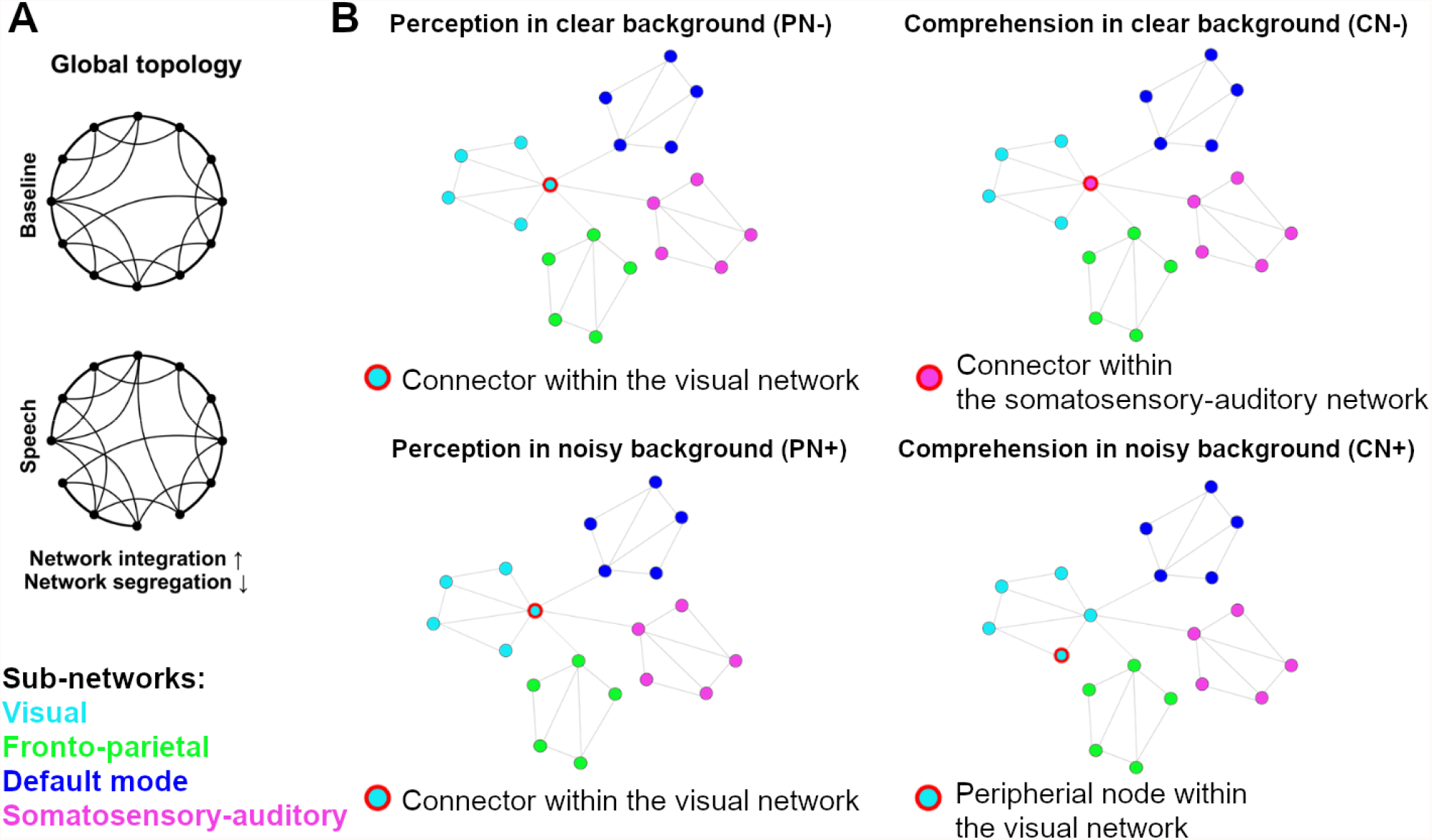
Conceptual illustrations of the main findings. **A**. Global topology observed during baseline (passive perception of bubble noise) and speech processing. Compared to the baseline, speech processing led to increased network integration and reduced network segregation, indicating an overall increase of information exchange across the whole brain. **B**. The left-vOT’s sub-network and its functional role in the sub-network. The area adaptively changed its functional role and sub-network depending on the speech processing situation. Red circle = Left-vOT node, cyan = visual network, green = fronto-parietal network, dark blue = default mode network, magenta = sensorimotor-auditory network.

At *the global level*, we found that, compared to passive exposure to unintelligible conversation noises, processing intelligible speech led to a more integrated and less segregated network. This evidence of an overall increase of information exchange across the whole brain is coherent with existing observations that processing speech recruits highly distributed brain areas that are involved in the analyses of acoustic, phonological, semantic, and syntactic information (Friederici, 2002; Vigneau et al., 2006; Cooke et al., 2006; Price, 2012; Huth et al., 2016; Abrams et al., 2020). The reorganization of the global network toward distributed processing observed here could be accounted for by the fact that, in the present protocol, participants were required to process entire sentences either to decide whether the same sentence was presented twice in a row or to extract their meanings. A similar global network reorganization has indeed been reported in other studies involving sentence reading and comprehension (Zhu et al., 2013; Feng et al., 2015). Importantly, as mentioned in the Results section, the fact that there was no difference between the four speech processing conditions at the global level allowed us to directly examine the role of the left-vOT between these conditions at the sub-network and nodal levels under the context of a similar global network organization.

More subtle network reconfigurations were revealed by the analyses at the sub-network level. The analyses of community structures showed a similar partition of sub-networks across the different speech processing conditions as well as in the baseline condition. These sub-networks consist of fronto-parietal network, default mode network, visual network, and sensorimotor-auditory network. The stable partition of sub-networks across the different processing contexts might reflect a common meso-scale organization of the brain network (Betzel 2020) engaged by both speech and non-verbal auditory processing (Price et al., 2005). The fronto-parietal network, default mode network and visual network are canonical sub-networks that have been consistently revealed by previous studies using functional connectivity (Greicius et al., 2003; Fox et al., 2005; Damoiseaux et al., 2006; Smith et al., 2009; Power et al., 2011; Laird et al., 2011; Cole et al., 2014). The fronto-parietal network mainly covers lateral prefrontal cortex, middle/inferior frontal regions, and posterior parietal cortex, and is regarded as a core system involved in a variety of tasks by implementing cognitive control and executive functions (Dosenbach et al., 2008; Cole et al., 2013). The default mode network is associated with various internal cognitive functions such as episodic memory and self-reference (Spreng et al., 2009; Davey et al., 2016), and mainly covers regions that are activated in task-free contexts (Raichle et al. 2001; Greicius et al. 2003), including medial prefrontal cortex, precuneus, and posterior cingulate regions. The visual network corresponds to the visual processing system that comprises visual areas. It is also consistently revealed in studies using both resting-state and tasks (Smith et al., 2009; Power et al., 2011; Cole et al., 2014). On the contrary, the sensorimotor-auditory network, which covers auditory and sensorimotor cortex, is more specifically considered as the main sub-network for auditory and speech processing, which is in line with the findings that sensorimotor cortex is also involved in speech perception and comprehension (Ito et al., 2009; D’Ausilio et al., 2009; Pulvermuller et al., 2010; Londei et al., 2010; D’Ausilio et al., 2012; Cogan et al., 2014; Schomers et al., 2016).

Importantly, despite the consistency of the sub-network partition across conditions, the analyses focusing on the left-vOT showed that the sub-network to which the left-vOT belonged varied depending on task demands and the clarity of speech signal. The left-vOT affiliated with the fronto-parietal network in the baseline where unintelligible auditory noises were presented without any task demands. This result is in line with previous studies that showed strong intrinsic connectivity between the left-vOT and fronto-parietal regions (Vogel et al., 2012; Chen et al., 2019). Interestingly, using Independent Component Analysis on intrinsic activity extracted from an auditory lexical decision task, López-Barroso et al. (2020) also found that the left-vOT is the only region that belonged to both the left fronto-parietal network and the lateral visual network. These findings, together with our results, suggest that the left-vOT could be a part of the fronto-parietal system in task-free situations such as resting-state or passive exposure to noise, while it might also maintain a subtle link with the visual system (López-Barroso et al., 2020). Here, we showed that this subtle link became obvious when spoken sentences were presented in a low-level perception task regardless of the clarity of the speech signal. In this case, the left-vOT belonged to the visual network (**Fig. 2**) and the investigation of its role at the nodal level showed that it interacted with the other sub-networks at the highest level among the other brain regions as reflected in its high values of participation coefficient (**Fig. 3A**). This observation indicates that, when one listens to speech, the left-vOT acts as a connector that links the visual system and the other subsystems. The results of *degree* and *clustering* also supported this view by showing that the left-vOT tended to connect with distributed regions rather than joining in a local cluster (i.e., low *clustering*), although it is unlikely to be a degree-based hub (i.e., intermediate *degree*). Together, these results provide empirical evidence in favor of the claim of the Interactive Account that the left-vOT acts as an interface between the visual system and high-order systems, and it receives various types of representations and information from high-order systems in a top-down manner.

Nevertheless, a more nuanced interpretation of the role of the left-vOT as an interface between the visual system and other high-order systems is called for if we consider the pattern of connectivity in the sentence comprehension task. The analysis conducted at both the sub-network and nodal levels revealed that extracting meaning from spoken sentences modified the functional status of the left-vOT compared to the speech perception task. Moreover, as discussed below, such modification of the functional status also depended on the clarity of the speech signal.

Firstly, when the sentences were clearly audible (no background noise), the left-vOT still held the highest level of communication with the different sub-networks as illustrated by the rank of the *participation coefficient* of this area compared to the other nodes (**Fig. 3A:** CN-). However, instead of belonging to the visual network, the area became part of the sensorimotor-auditory network that is typically recruited during spoken language processing (**Fig. 2**). As reported in the literature, the involvement of this network generally increases when speech comprehension is explicitly required (Hickok et al., 2007; Friederici et al., 2012; Schomers et al., 2016). Our observation that during spoken sentence comprehension the left-vOT became part of this network is coherent with previous observations. The left-vOT reported here was functionally localized using a word reading task. The extracted ROI was centered at x = -47, y = -55, z = -17 (MNI), which corresponds the location of *classical Visual Word Form Area* (cVWFA) or the middle occipito-temporal sulcus identified by Lerma-Usabiaga et al. (2018). The authors reported that the cVWFA is anatomically connected with the angular gyrus and the inferior frontal gyrus and is responsible for integration of linguistic information with the spoken language network. Additionally, our finding that the left-vOT could belong to the spoken language system under some circumstances is also in line with Saygin et al.’s longitudinal data (2016) showing that, even before the written language is acquired, the future location of what will become the VWFA is determined by its pattern of anatomical connectivity with the spoken language system. Here, we provided additional evidence that, at least in adults, the possibility that the area would become part of the spoken language network is even more likely when high-order information processing, such as speech comprehension, is required.

Secondly, the pattern of left-vOT connectivity changed again when the sentences were presented against background noise: The left-vOT no longer belonged to the sensorimotor-auditory network but returned to the visual network, and its functional role turned from a connector into a simple peripheral node. It is worth noting that in this specific condition, the task performance decreased significantly both in terms of accuracy score and processing speed, which indicates an increase of task difficulty and cognitive demands when one had to comprehend speech in a noisy environment (Peelle et al., 2018). Related to this explanation, we also observed, in the CN+ condition, a negative correlation between the reaction times and the participation coefficient. Participants who showed less difficulties in comprehending the sentences despite the background noise were also those who showed higher communication between the left-vOT and the other sub-networks. This inter-individual variability suggests that, while the left-vOT plays a role as peripheral node at the group level, its capacity to communicate with other sub-networks could predict participants’ task performance especially in challenging speech processing situations. The disengagement of the left-vOT from the spoken language system, as reflected by the change of sub-network (from the sensorimotor-auditory network to visual network) and functional role (from connector to peripheral node) in the most difficult speech comprehension situation, mirrors some previous findings that the degree of activation of some higher-order areas in the language network does not always increase with task difficulties (Obleser et al., 2007; Vagharchakian et al., 2012). For instance, Obleser et al. (2007) used a speech perception task in which they manipulated the clarity (S/N ratio) and the semantic predictability of speech signal and found that the benefit of semantic predictability was strongest at an intermediate level of speech degradation. The improvement in comprehension of degraded speech thanks to the semantic predictability was associated with an increase in activity and functional connectivity between higher-order cortical areas. However, such benefits disappeared when the signal was severely degraded.

In sum, in addition to the consensus on the central role of the left-vOT in reading, the brain connectivity findings reported here further demonstrates that this area also plays an adaptive and pivotal role in the communication between different brain systems during speech processing. These findings are in line with the Interactive Account which considers that, beyond its key role in reading, the left-vOT is an interface between bottom-up sensory inputs and top-down predictions that call on non-visual stimulus attributes (Price and Devlin 2011). This role could be supported by the left-vOT’s intrinsic and anatomical connections with various brain regions and systems (Koyama et al., 2010; Vogel et al., 2012; Yeatman et al., 2013; Bouhali et al., 2014; Saygin et al., 2016; Stevens et al., 2017; Chen et al., 2019; Li et al., 2020), and, as shown here, is modulated by the interaction between bottom-up (clarity of the signal) and top-down (task demands) factors. Specifically, in a basic speech perceptual task, the area belonged to the visual system and contributed to the exchange of information between the visual and the other systems. Most probably due to its location at the transition between the occipital and the temporal cortex, in a higher-level speech comprehension task, the area became part of the spoken language network and continued to participate in the communication with the other systems (Lerma-Usabiaga et al., 2018). However, when speech comprehension could no longer be performed at a satisfactory level due to a suboptimal listening situation, the left-vOT disengaged from the spoken language network and turned back to the visual system. At the same time, it no longer acted as a connector between different processing systems.

Although several studies have reported the activation of the left-vOT during speech processing (Booth et al., 2002; Cao et al., 2010; Ludersdorfer et al., 2015, 2016), our study is the first to reveal the functional role of this “reading area” in speech processing beyond its widely accepted contribution to phonology-to-orthography mapping process (Dehaene and Cohen, 2011). It shows that the left-vOT adapts its function to support the communication between the visual system and other sub-networks according to task demands and the characteristic of sensory input. These varying patterns of functional connectivity provide a valid mechanism that explains the on-going debate on the role of the left-vOT in different processing contexts (Dehaene and Cohen, 2011; Devlin and Price, 2011) and is in line with an increasing number of studies that show the implication of the area in large-scale brain networks (Bouhali et al., 2014; Saygin et al., 2016; Chen et al., 2019). At the theoretical level, our finding provides empirical evidence in favor of a key assumption of the Interactive Account that the left-vOT acts as a convergent zone having multiple functions depending on the interactions with other cortical or subcortical areas (Damasio & Damasio, 1994; Price & Devlin, 2003). More studies are nevertheless needed to further explore this observation, for instance, whether the adaptive role of the left-vOT is supported by a single population of neurons that adjusts its pattern of connectivity according to the processing context, or by different subpopulations of neurons located in the same area that may have different pattern of connectivity, as suggested by Pattamadilok et al. (2019, see also Prince & Devlin 2003). Also, using measures of neural propagation with higher temporal-resolution and a causal interventional approach (Rogasch et al., 2013; Bortoletto et al., 2015) could help to clarify the temporal dynamics, the direction of the functional connectivity between the left-vOT and the other brain areas, as well as the causal relationship between the pattern of brain connectivity and language processing performance.

## Acknowledgements

This work was supported by the French Ministry of Research: ANR-13-JSH2-0002 and ANR-19-CE28-0001-01 (to C.P.), ANR-16-CONV-0002 (ILCB), ANR-11-LABX-0036 (BLRI) and the Excellence Initiative of Aix-Marseille University (A*MIDEX). It was performed in the Center IRM-INT (UMR 7289, AMU-CNRS), platform member of France Life Imaging network (grant ANR-11-INBS-0006). We warmly thank Dr. Agnès Trébuchon for taking medical responsibility during the study. Center de Calcul Intensif d’Aix-Marseille is acknowledged for granting access to its high-performance computing resources.

