## Supplementary_Information for "Graph theoretical analysis reveals the adaptive role of the left ventral occipito-temporal cortex in the brain networks during speech processing"

Methods

***A) Group left-vOT defined by the visual localizer task***

The pre-processed functional data obtained in the visual localizer task were fitted into a general linear model (GLM) with a block-design using AFNI. The same set of nuisance regressors as the spoken sentence processing tasks were included, that are 1) 24 head motion regressors: the six motion parameters, their temporal derivatives, and all their corresponding squared time series and, 2) 26 physiological regressors: the mean time-series and the first twelve principal components of white matter and of CSF which were extracted by using the aCompCor method (Behzadi et al., 2007), as well as the cosine-basis regressors estimated by fMRIPrep. Motion contaminated volumes were identified by using framewise displacement (FD) and were censored along with the prior volume if their FD > 0.5mm. On average, 1.8% of the volumes were censored for the visual localizer task. The group level analysis was then performed to compute the contrast *words - consonant strings* by using pairwise t-test in AFNI (3dttest++). To combine the group left-vOT with the set of 263 spherical ROIs that are described in the main text (cf. Power et al. 2011), the group left-vOT with the same volume of 81 voxels as a spherical ROI was extracted. To this aim, the group activation map of the contrast *words - consonant strings* was thresholded to extract the strongest cluster with a volume of 81 voxels by adjusting the significance threshold (T > 5.2536, *p* < 2.5e-5). The spatial distribution of the final set of ROIs is shown in **Fig. S1A**.

***B) Network construction of the symmetrical ROIs set (Di et al., 2014)***

A symmetrical set of 402 ROIs were taken from (Di et al., 2014) to confirm the global and nodal results reported in the main text while controlling the spatial bias in lateralization. The same procedure as described in the main text (see Network Construction) was performed on this set of ROIs. Specifically, the 402 ROIs with a radius of 5 mm were intersected with the group-averaged gray matter mask to exclude the ROIs that are outside the gray matter, resulting in 392 ROIs. The 10 ROIs that were removed were mainly subcortical nuclei located outside the gray matter mask. Additionally, the left-vOT identified in the visual localizer task and its homogeneous region in the right hemisphere were included into the symmetrical ROIs set. Finally, three ROIs in the left Fusiform gyrus and three ROIs in right Fusiform gyrus were further removed due to the overlap with functionally localized left-vOT and its homogeneous region, respectively. Altogether, this resulted in 388 ROIs in the symmetrical set (see **Fig. S1B** for the spatial distribution of this set of ROIs). The beta-series connectivity and Fisher-z transformation were then applied on the symmetrical ROIs set to construct a brain network for each condition and each participant. The density range here was identified between 14% and 18% at steps of 1%. The lower bound, 14% density, is the sparsest density at which 90% of all the networks are fully-connected, without any significant between-condition difference in the size of the LCC. The upper bound, 18% density, is the sparsest density where all the networks are fully-connected networks.

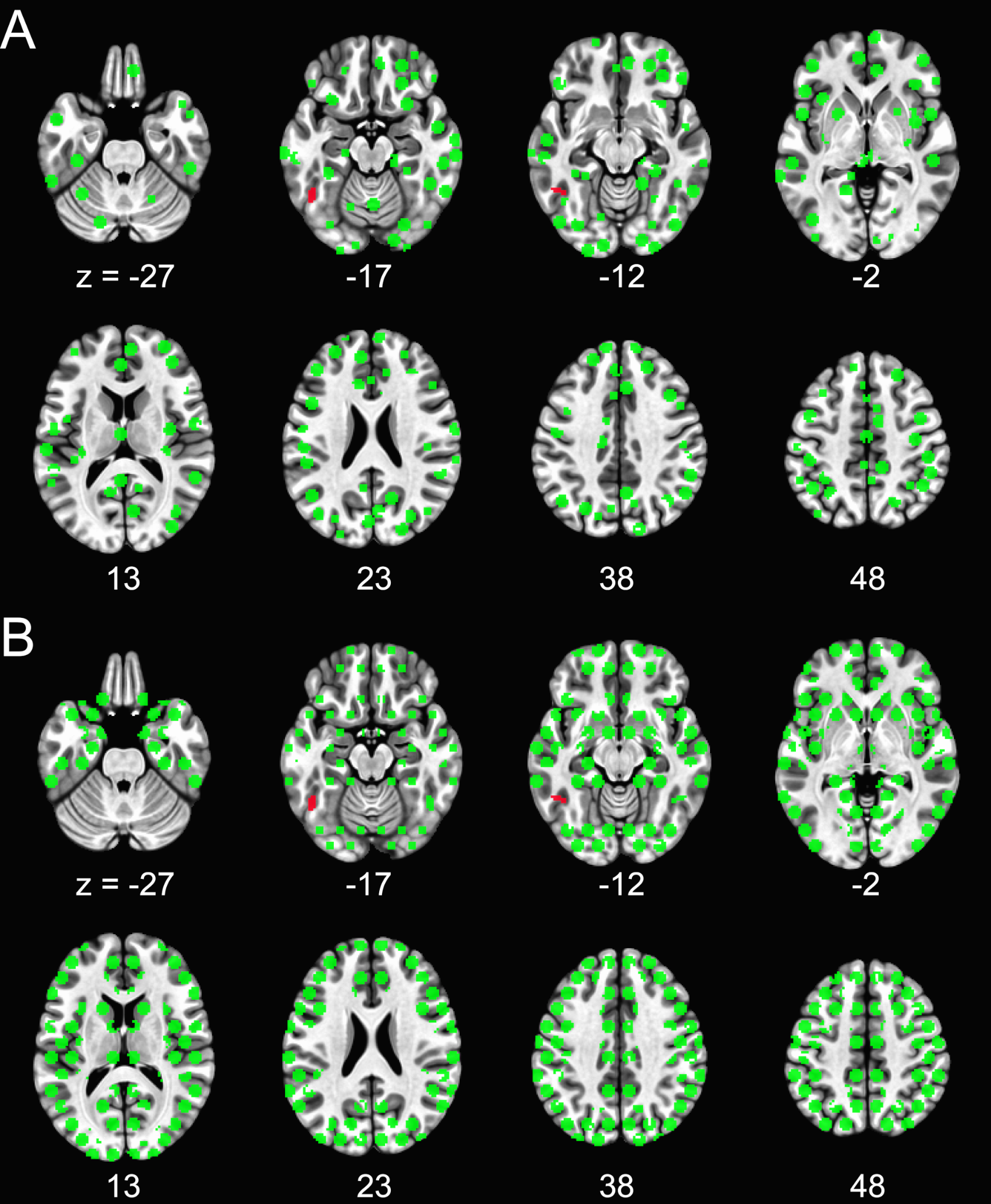

**Figure S1.** **A.** The spatial distribution of the set of 263 ROIs originally taken from Power et al. (2011). **B.** The spatial distribution of the set of 388 ROIs originally taken from Di et al. (2014). *Notes:* Each ROI is a sphere with a radius of 5mm (i.e., 81 voxels) and is intersected with the group-averaged gray matter mask to exclude areas that are outside the gray matter. The group left-vOT with 81 voxels (colored in red) is centered at MNI x = -47, y = -55, z = -17.

***C) Sub-network labeling***

The analysis of community detection identified four sub-networks for both baseline and speech processing conditions. As mentioned in the Results section of the main text, the community structures were similar across conditions. We labeled these four sub-networks as visual network, fronto-parietal network, default mode network, and sensorimotor-auditory network based on the original community structure and labels defined by Power et al. (2011). As illustrated in **Figure S2**, the cyan and dark blue sub-networks highly matched the “visual network” and “default mode network” reported in Power et al. (2011), respectively. The green sub-network largely overlapped with the fronto-parietal task control, ventral and dorsal attention, and salience networks in Power et al. (2011) and was labeled “fronto-parietal network” in our study. Finally, the magenta sub-network contained two somatosensory-motor networks (month and hand) and the auditory network reported in Power et al. (2011). Although it also overlapped with several subcortical networks (i.e., cingulo-opercular task control, subcortical, cerebellar), we labeled this sub-network as “Sensorimotor-auditory Network”.

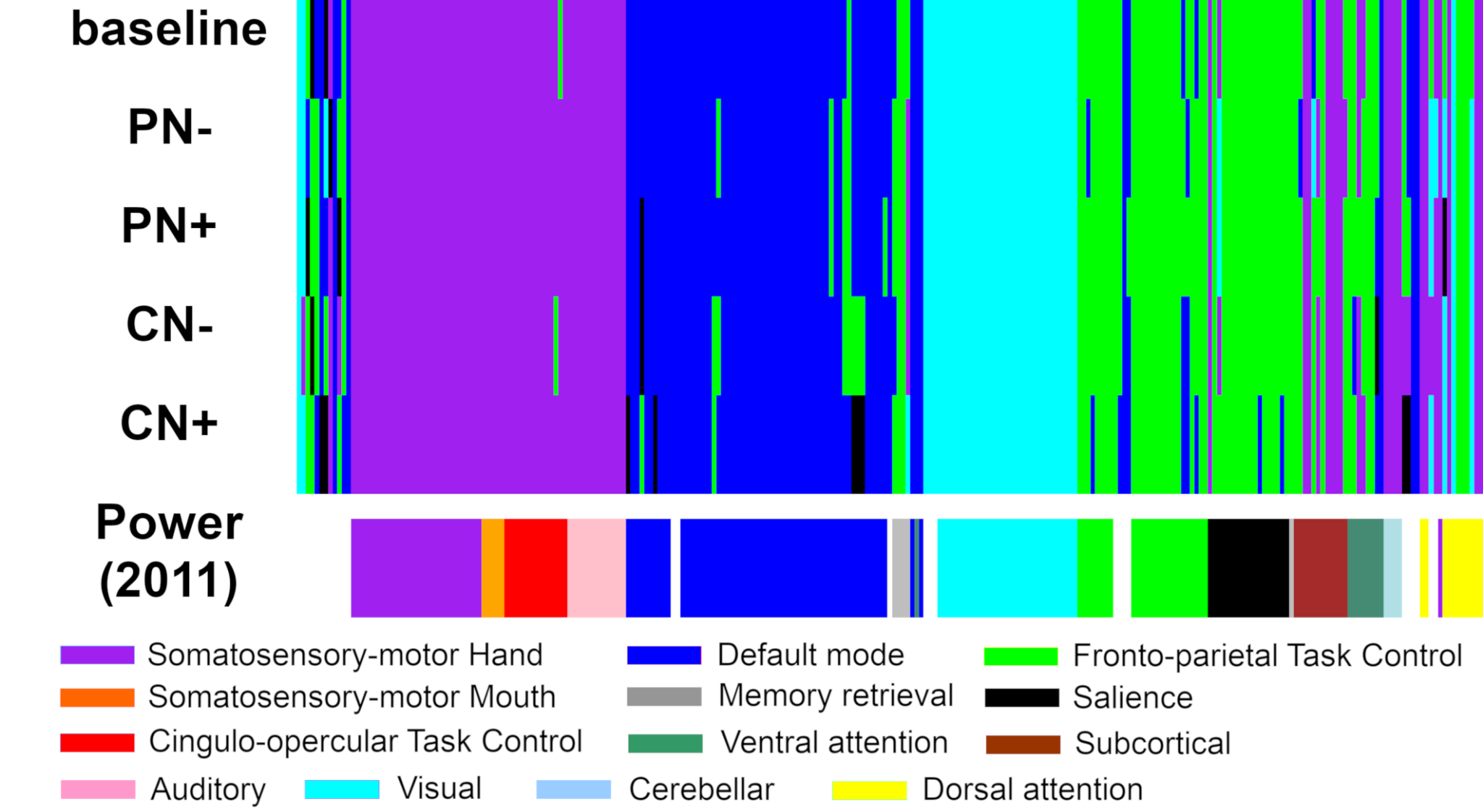

**Figure S2.** Community structures (partitions) and labeling. The top five rows show the community structures identified for baseline and speech processing conditions, which mainly contain four sub-networks that are colored cyan (visual network), green (fronto-parietal network), dark blue (default mode network) and magenta (sensorimotor-auditory network). The bottom row shows the community structure defined by Power et al. (2011) through meta-analysis, where white bars present ROIs that cannot be clearly assigned into a sub-network. The legend shows the labels originally defined by Power et al. (2011).

Results

To ascertain that the results reported in the main document, where the analyses was conducted on the ROIs from Power et al. (2011), were not restricted to a specific set of ROIs or to a single density (15%), we conducted additional analyses: 1) across the range of densities 15%-22% with the FDR correction using Power et al.’s ROI (2011) and 2) using the symmetrical ROIs set across the range of densities 14%-18% with the FDR correction. As respectively shown in **Tables S1** and **S2**, the new analyses confirmed the patterns of functional connectivity reported in the main document.

**Table S1.** Between-conditions significance in global and nodal metrics of brain networks based on the 263 ROIs (Power et al., 2011). The results were corrected with FDR across the range of densities (15%-22%). FDR corrected ps < 0.05 are highlighted in bold.

| Density | | | | | | | | |
| --- | --- | --- | --- | --- | --- | --- | --- | --- |
|  | **15%** | **16%** | **17%** | **18%** | **19%** | **20%** | **21%** | **22%** |
| clustering coefficient | **0.009** | **0.009** | **0.009** | **0.009** | **0.009** | **0.009** | **0.009** | **0.009** |
| global efficiency | **0.005** | **0.005** | **0.005** | **0.005** | **0.005** | **0.005** | **0.005** | **0.005** |
| number of communities | 0.391 | 0.432 | 0.432 | 0.432 | 0.432 | 0.432 | 0.391 | 0.432 |
| modularity Q | 0.426 | 0.426 | 0.426 | 0.426 | 0.426 | 0.426 | 0.426 | 0.426 |
| degree | 0.058 | 0.058 | 0.058 | 0.058 | 0.058 | 0.058 | 0.058 | 0.058 |
| clustering (nodal) | 0.670 | 0.670 | 0.670 | 0.670 | 0.670 | 0.670 | 0.670 | 0.670 |
| within-module z-score | 0.187 | 0.187 | 0.187 | 0.187 | 0.229 | 0.279 | 0.267 | 0.267 |
| participation coefficient | **0.002** | **0.002** | **0.002** | **0.002** | **0.004** | **0.028** | **0.028** | 0.074 |

**Table S2.** Between-conditions significance in global and nodal metrics of brain networks based on the 388 ROIs (Di et al., 2014). The results were corrected with FDR across the range of densities (14%-18%). FDR corrected ps < 0.05 are highlighted in bold.

| Density | | | | | |
| --- | --- | --- | --- | --- | --- |
|  | **14%** | **15%** | **16%** | **17%** | **18%** |
| clustering coefficient | **0.003** | **0.003** | **0.003** | **0.003** | **0.003** |
| global efficiency | **0.003** | **0.003** | **0.003** | **0.003** | **0.003** |
| number of communities | 0.105 | 0.105 | 0.105 | 0.105 | 0.148 |
| modularity Q | 0.795 | 0.795 | 0.795 | 0.795 | 0.795 |
| degree | **0.024** | **0.024** | **0.024** | **0.024** | **0.024** |
| clustering (nodal) | 0.577 | 0.416 | 0.416 | 0.416 | 0.416 |
| within-module z-score | 0.147 | 0.147 | 0.092 | 0.092 | 0.112 |
| participation coefficient | **0.050** | **0.046** | **0.046** | **0.046** | **0.050** |

At the nodal level, the analysis of the functional role of the left-vOT conducted on the symmetrical ROIs set at the density of 15% also confirmed the results reported in the main document. Among the 388 nodes, the left-vOT node was ranked 51^st^ (PN-), 20^th^ (PN+), 165^th^ (CN-), and 59^th^ (CN+) in order of *degree*, while the left-vOT node was ranked 360^th^ (PN-), 336^th^ (PN+), 349^th^ (CN-), and 280^th^ (CN+) in order of *clustering*. The four spoken sentence conditions differed in *degree* (*p* < 0.019) but not in *clustering* (*p* > 0.23).

**Fig. S3A** showed the rank of vOT in terms of *participation coefficient* and *within-module z-score*. The left-vOT node was ranked 1^st^ (PN-), 2^nd^ (PN+), 12^th^ (CN-), and 78^th^ (CN+) in order of *participation coefficient*, while the left-vOT node was ranked 169^th^ (PN-), 195^th^ (PN+), 259^th^ (CN-), and 197^th^ (CN+) in order of *within-module z-score*. The comparison in *participation coefficient* showed a significant difference between conditions (*p* < 0.019), while no significant difference was found in *within-module z-score* (*p* > 0.12). The *post-hoc* tests confirmed that the *participation coefficient* in the CN+ condition was significantly lower than in the PN- (*p* < 0.0035) and the PN+ (*p* < 0.0081) conditions, but not the CN- condition (**Fig. S3B**). As shown in **Fig. S3C**, the correlation analysis showed that *participation coefficient* and reaction time was negatively correlated in the CN+ condition (Pearson’s r = -0.48, *p* < 0.019), while the correlations were not significant in other conditions (all *p*s > 0.31).

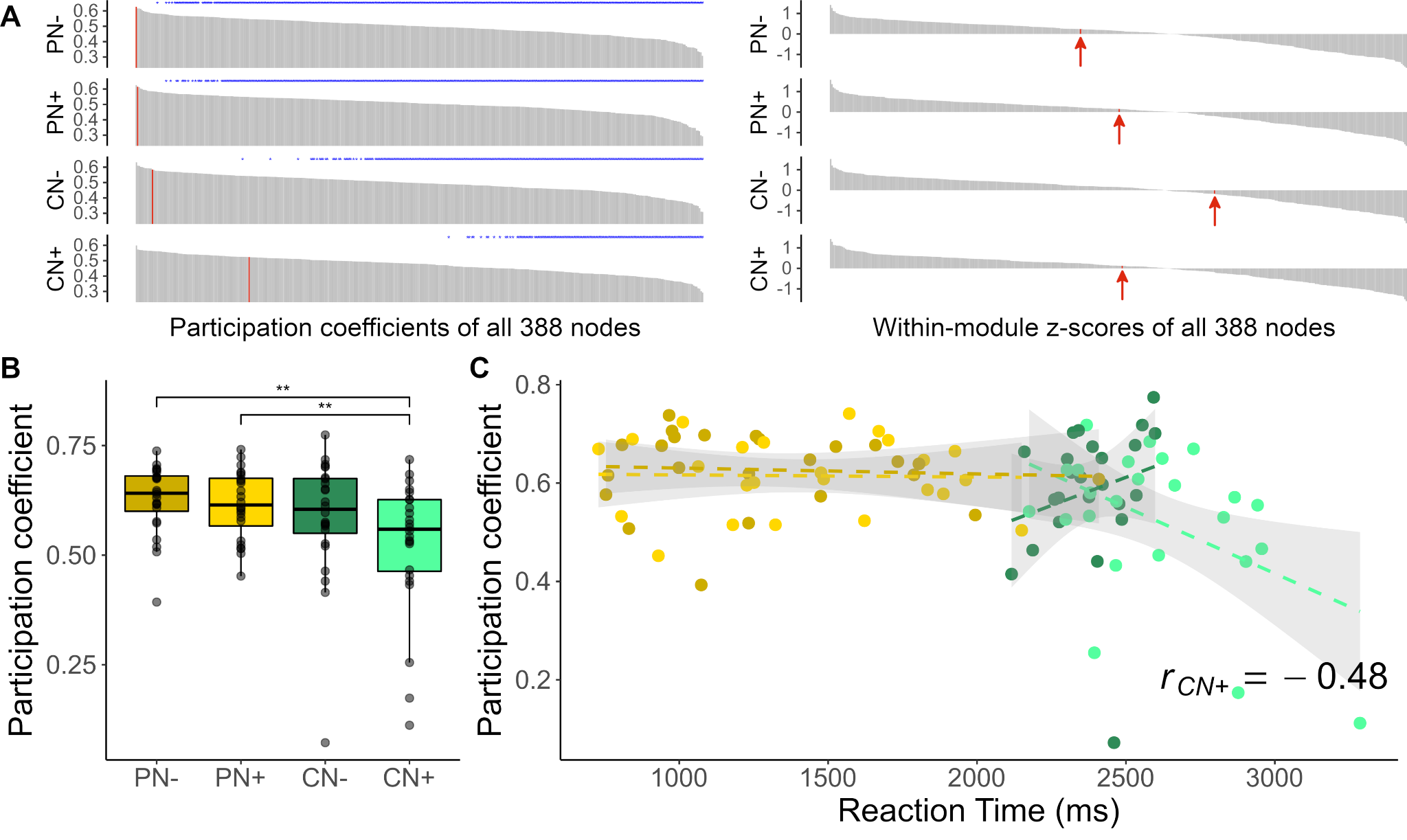

**Figure S3.** **A.** The left-vOT node showed high *participation coefficient* and intermediate *within-module z-score* among all the 388 nodes. The blue asterisks indicate the nodes that showed significantly lower *participation coefficients* than the left-vOT (paired permutation test, *p* < 0.05 unc.). **B.** The *participation coefficient* of the left-vOT node was significantly lower in the CN+ condition than in the PN- and PN+ conditions, but not the CN- condition. **C.** The *participation coefficient* and reaction time was negatively correlated in the CN+ condition (light green dots and dashed line; Pearson’s r = -0.48, *p* < 0.019), while the correlations are not significant in other conditions (all *p*s > 0.31).
